# Paleogene mass extinction and ongoing Neogene recolonization shape the tropical African flora

**DOI:** 10.1101/2025.04.01.646671

**Authors:** Daniela Mellado-Mansilla, Anna Weigand, Patrick Weigelt, Holger Kreft, Michael Kessler

## Abstract

Tropical regions are known to host most of the global biodiversity, yet species richness varies drastically between continents. Tropical Africa has long been recognized as the “odd man out”, exhibiting lower plant diversity compared to other tropical continents, but the underlying causes remain debated. Here, we use ferns as a model group to explore the processes responsible for the low diversity of African plants. We find that the current fern diversity in Africa, particularly in humid regions, is up to 84% lower than under similar climatic conditions in the Americas and Asia. Unlike on these continents, where 55–60% of extant fern diversity is the result of in-situ diversification of Gondwanan lineages, only 16% of African fern diversity originates from such lineages. This discrepancy points to significant African extinction periods during the Paleogene and mid-Miocene, likely driven by elevated temperatures and aridification. In contrast, 54% of the extant African fern diversity can be attributed to approximately 530 intercontinental dispersal events during the Neogene, indicating ongoing recolonization of Africa. Our findings provide unprecedented insight into the evolutionary forces shaping plant diversity in tropical Africa and highlight the potential risks posed by ongoing climate change to its botanical heritage.

## Main

Tropical plant diversity is unevenly distributed across continents. Tropical Africa, the “odd man out”^1^, stands out among the tropical regions by having notably lower species richness and phylogenetic diversity compared to tropical America and Asia^2,3^. This disparity spans numerous plant groups, including palms^4^, epiphytes^5^, and other emblematic tropical taxa. Estimates suggest that Africa hosts around 56,500 vascular plant species, far fewer than tropical America (118,308) and Asia (50,000)^6–8^. Similarly, Africa has only about 6,000 tree species, compared to approximately 25,000 in both America and Asia^9^. Among ferns, this anomaly is even more pronounced^10^: continental Africa harbors about 1,400 species^11^ compared to 3,360 in tropical America^12^ and 4,500 in tropical Asia^13^.

Four leading hypotheses have been proposed to explain this intercontinental disparity^3,14^. First, the “climate-suitability” hypothesis proposes that current climate in Africa is less suitable for plant growth than those of America or Asia^15^. Second, a variant of this idea, which we term the “habitat-extent” hypothesis, suggests that the limited current extent of suitable humid habitats in Africa restricts habitat diversity by constraining speciation rates and increasing extinction risks due to smaller population sizes^10,16^. Third, the “climate-stability” hypothesis, argues that America and Asia experienced less dramatic climatic shifts than Africa throughout their geological histories, particularly in precipitation patterns, leading to habitat loss and higher extinction rates in Africa^17^. Fourth and finally, the “time-to-speciation” hypothesis and the associated “time-integrated species-area effect” suggest that ancient lineages in Asia and America had more time to diversify in large, stable habitats^18^. These not mutually exclusive hypotheses reflecting different aspects of fundamental intercontinental differences, have not yet been widely tested in plants and remain unexplored for African ferns.

With 12,000 accepted species^19^, ferns and lycophytes are an ideal model group to test these hypotheses. With a deep evolutionary history spanning over 400 million years^20^, ferns predate the break-up of Pangea and Gondwana, and dominated terrestrial vegetation along with gymnosperms until the rise of the angiosperms to ecological dominance in the Cretaceous^21^. Ferns later diversified into approximately 80% of extant taxa by exploiting new niches created during the angiosperm dominance, becoming a characteristic component of humid tropical forests^22,23^. Fern diversity is highly constrained by their sensitivity to water stress^10,24^, making them potential indicators of past climatic shifts. Additionally, their small, wind-dispersed spores, can travel thousands of kilometers^25,26^, leading to numerous intercontinental dispersal events^27^. Since ferns do not depend on animals as pollinators and dispersers, their distributions tend to be more in balance with environmental conditions than those of seed plants^28^.

Here, we aimed to understand the underlying mechanisms causing the low plant diversity in tropical Africa compared to other tropical regions using ferns as a model group. Using a phylogeny including roughly half of the global fern species^29^ and a global model of fern species richness^30^, our findings suggests that the low diversity of tropical African ferns cannot be explained by current environmental conditions but rather declined during massive extinctions in arid periods until the late Miocene, followed by ongoing recolonization via intercontinental dispersal.

## Results

### Macroecological diversity patterns

Our spatial model of fern diversity was based on Weigand et al. (2020), who used 1,243 regional floristic inventories globally and included eight climatic variables plus the area of the survey units, to calculate an environment-based statistical model of fern richness for a fixed grid cell size of 7,666 km^2^ that accounted for 68.9% of the variance (Supplementary Table 1). Introducing continents as an additional factor increased the model’s explanatory power by 5.9%, revealing significant intercontinental differences in fern diversity beyond modern environmental differences (Fig. 1A, Supplementary Table 1). Tropical Asia (estimate = 0.66) and South America (0.41) were modeled to be more species-rich compared to Africa, which had an estimate of 0 (Supplementary Table 1).

**Figure 1.**
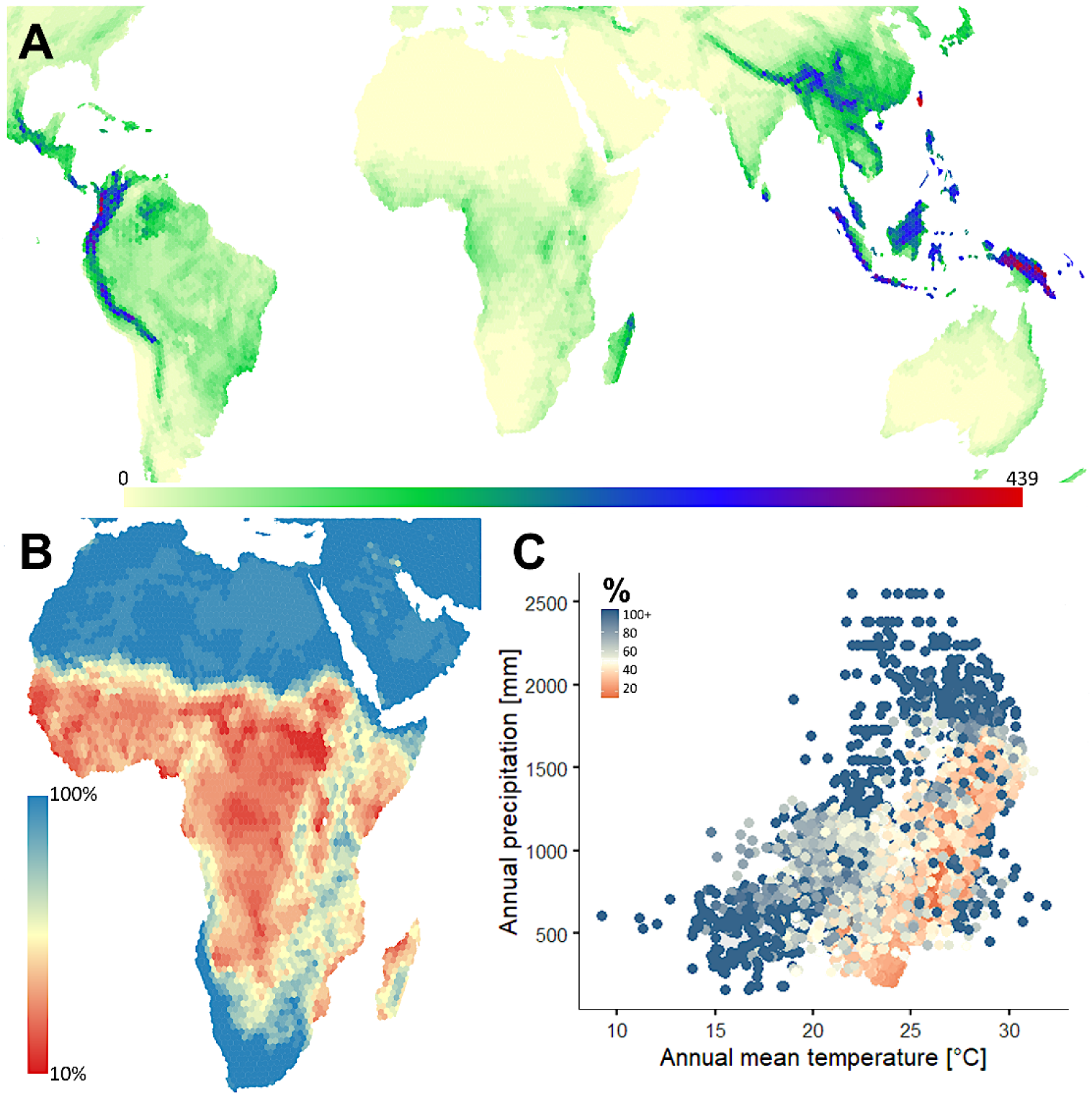
A. Modelled regional fern richness at 7,666 km^2^ grid cell resolution, showing low species richness in Africa and high richness in tropical America and Asia^30^. B. Percentage of missing fern species richness in Africa based on the comparison of models based on non-African with only-African environment-richness relationships. C. Relationship of percentage of missing fern species richness (color gradient) in relation to annual mean temperature and annual precipitation.

To further investigate the discrepancy in African fern diversity, we used climate-richness models based only on tropical America and Asia to predict potential fern richness in Africa, assuming similar climate-diversity relationships. By subtracting this from the actual richness in Africa, this comparison revealed that Africa has on average, 34.3% fewer fern species per grid-cell than would be expected based on climate alone (Fig. 1B). This deficit was especially pronounced in humid and hot regions, where Africa’s fern diversity was up to 84.4% lower than predicted. This pattern contradicts the “climate-suitability” hypothesis, which proposes that current climatic conditions explain inter-continental differences in plant diversity ^15^. It also shows that African rainforests have a much higher deficiency of fern richness than arid ecosystems, suggesting that the causes for low African fern diversity may be related to past precipitation regimes ^31^, as proposed by the “climate-stability” hypothesis.

We also tested the “habitat-extent” hypothesis by incorporating the extent of cloud forest habitats ^32^—one of the most species-rich environments for ferns ^22,30,33^—into the model. While the inclusion of this variable slightly improved the model’s explanatory power slightly from 74.4% to 74.8% and decreased the dAIC from 48501 to 48091, it did not replace the continental factor, reinforcing the importance of biogeographic history in shaping African fern diversity.

### Macroevolution and the temporal buildup of tropical fern diversity

We continued by analyzing the evolutionary history of fern diversity in Africa which reveals a distinct pattern compared to tropical America and Asia (Fig. 2). For this analysis, we considered the entire evolutionary history of ferns, from 421 mya to the present, but defined ancient lineages as those that predate the K/T boundary (∼66 mya) because phylogenetic inferences before that time may be challenging, as the K/T boundary was associated with a major extinction event among terrestrial biota, including ferns. Furthermore, the breakup of Gondwanan took place until about 70 mya, so that continental landmasses were still so close to each other that they were not effectively isolated for spore-dispersed plants such as ferns^3,27^.

**Figure 2.**
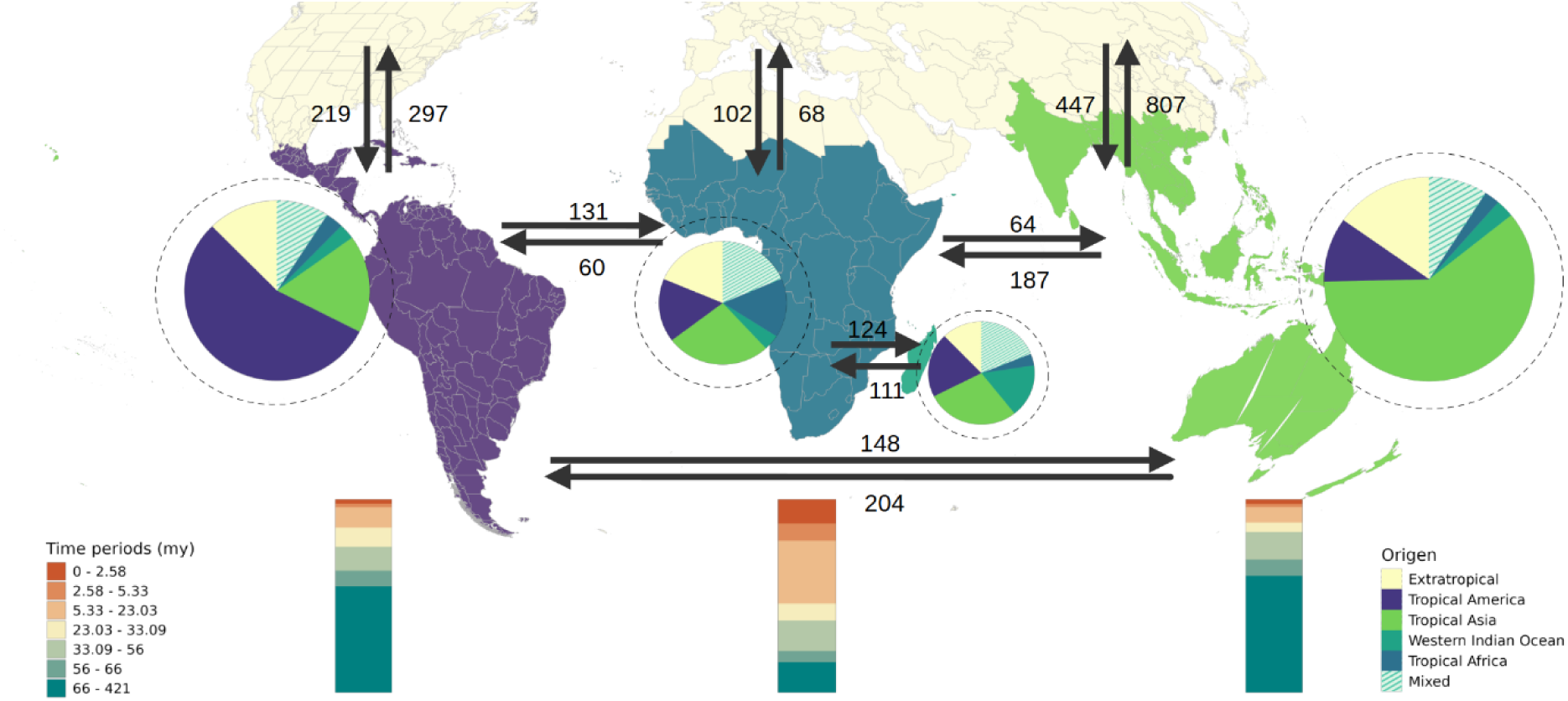
Summary graph of the dispersal event reconstructions of the buildup of fern diversity on the main tropical continents. The circles show the total numbers of fern species in each continent (dashed circles) and the numbers of species included in the phylogeny (filled circle), with the colored pie segments showing the biogeographical origins of the current fern species, corresponding to lineages present in a continent over the last 66 my. The bar graphs indicate the timing of the first appearance of an extant fern lineage on each continent. Lines indicate the direction of the dispersal events between areas, and the numbers correspond to the mean numbers of dispersal events.

We found that in Africa only 16 ± 4% (mean ± sd) of fern species included in our analysis belong to ancient lineages that survived on the African continent since the end of the Mesozoic (including “mixed” lineages, for which more than one continent including Africa was reconstructed as ancestral range). In contrast, 55 ± 7% of the tropical American and 60 ± 8% of the tropical Asian species included in the analysis are derived from such ancient Gondwanan lineages. The remaining African fern diversity (84 ± 7%) resulted from an estimated 531 ± 2 dispersal events that took place within the last 66 million years and subsequent diversification. This colonization was especially pronounced in the Neogene, with 54 ± 4% of the extant species deriving from 381 ± 4 dispersal events into Africa that took place in the last 23 million years, with a considerable percentage (13 ± 2%) derived of very recent dispersal events during the Pleistocene. In contrast, in America and Asia, 45 ± 4% and 40 ± 4% of the current species diversity, respectively, derived from 540 ± 2 and 720 ± 3 Cenozoic dispersal events, with only 2 ± 0.4% and 2 ± 0.6% species deriving from Pleistocene dispersal events.

## Discussion

The lower plant diversity of tropical Africa compared to America and Asia has long puzzled ecologists. Our study provides strong evidence that the low diversity of African ferns cannot be attributed solely to current climatic conditions or habitat extent, as also shown for angiosperms^3^. Importantly, we find that African fern diversity is depauperate in humid tropical forests but is similar to that of America and Asia in arid and cooler areas. This pattern is mirrored by that of angiosperms, where African rainforests are also known to be relatively depauperate^3,6^, whereas arid ecosystems like the southern African karoo and Mediterranean-type fynbos are among the most diverse globally^34^. This shows that the history of humid tropical forests is key to understanding the diversity disparity between continents.

Our dispersal event reconstructions indicate that, contrary to America and Asia, most fern species in Africa today are not derived from lineages that have persisted since the Gondwanan break-up, but rather are the result of Cenozoic and especially Neogene dispersal events and subsequent diversification. This suggests that pre-Cenozoic climatic conditions were unsuitable for the long-term survival of many fern species, and that numerous lineages originally occurring in Africa went extinct. Unfortunately, the hypothesis of the original presence of many fern lineages in Africa and their subsequent extinction cannot be directly tested due to the paucity of fern fossils from Africa^35^. However, fossil records from South America, Australia, and Antarctica predating the Gondwana breakup around 145-70 mya, reveal that the flora of these continents included many lineages of ferns and other vascular plants occurring across Gondwana which today survive only on some of the resulting landmasses, reflecting Cenozoic extinctions on the other continents following their separation^27,36,37^. Considering the central position of Africa in Gondwana, it is likely that many of the fern lineages that persisted throughout the Cenozoic in America and Asia were also originally present in Africa, suggesting that the low African fern diversity stems from extinctions as a result of Cenozoic climatic fluctuations as posited by the “climate-stability” hypothesis.

We propose the following scenario for the evolution of fern diversity on tropical continents and the higher extinction in Africa. Before its breakup, Gondwana had a largely tropical flora that showed regional differentiation but nevertheless was fairly homogenous, resulting in similar floras on the separating continents^37^. Globally, tropical climates are thought to have been widespread until the Eocene-Oligocene transition, when the growth of polar ice caps resulted in cold water currents and increased rainfall seasonality^38,39^. Tropical regions shrank and forest habitats shifted polewards, but while South America and Asia had extensive landmasses extending north- and southwards, allowing for forest migration, in Africa the northern arid belt and the limited southward extent of landmasses reduced opportunities for latitudinal forest expansion^14^. Survival in tropical America and Asia was further facilitated by the presence of extensive mountain ranges with humid forests, such as in the Andes, Himalayas, and New Guinea, which provided large refugia during periods of hot and dry climates, as experienced during the mid-Miocene climate optimum^40^. In Africa, in contrast, the major mountain ranges of Ethiopia and southern Africa have quite dry climates and the tropical mountains are mainly young volcanoes, which provide limited refugia and lead to higher extinction rates during dry climatic conditions of plant species dependent on tropical forest habitats than in tropical America or Asia^41^. During the late Miocene, humid tropical regions on all continents expanded again, providing extensive habitat for ferns and allowing for successful intercontinental dispersal and colonization, followed by diversification, as found by us and previous studies on individual African fern genera^42,43^. In contrast, in America and Asia, many Gondwanan lineages survived and diversified in continuously suitable habitats throughout the Cenozoic, in line with the “time-for-speciation” hypothesis. Intercontinental dispersal leading to colonization also occurred on these continents but contributed much less to overall fern diversity due to the persistence of Gondwanan lineages.

Our results have several important implications for the management and conservation of biodiversity in Africa. First, since other plant groups, especially those inhabiting humid forest habitats, show similarly low diversity in Africa compared to America and Asia^2–4,14^, it is likely that the evolutionary and community assembly processes proposed here for ferns apply to a broader range of plant groups and potentially also to other groups of organisms. In fact, the high dispersal ability of ferns could have provided a significant advantage in recolonization in comparison to other plant groups with less effective long-distance dispersal. For instance, palms, which mostly have large, animal-dispersed seeds, only have 65 species in Africa, compared to 800 and 1200 in America and Asia, respectively^44^. Second, the late arrival of new fern lineages combined with the low current diversity suggests that African fern diversity may not have yet saturated and would continue accumulating species if left under natural conditions without human impact. However, depauperate species assemblages may be less resilient to environmental changes and more sensitive to species losses than assemblages with functionally redundant species^45,46^. Third, physiological drought stress in plants is not solely determined by water input via precipitation, but is also influenced by water loss via evapotranspiration, a temperature-dependent process^47^. Indeed, high temperatures have been identified as the primary drivers of global fern extinctions over evolutionary time scales^48^, which is in accordance with our finding of high extinction resulting from hot and arid periods up into the Miocene. This raises concerns about the future of African fern diversity under climate change. Predictions of higher temperatures and stronger aridity in Africa, akin to the mid-Miocene climate optimum, may lead to a new plant extinction crisis on the continent, with profound implications for ecosystem functioning and human well-being. This potential scenario could be further exacerbated by the widespread destruction of African tropical rainforests^49^, not only in the lowlands, but most crucially in the mountains, where they would act as climate change refugia^50^. Our study underscores the crucial importance of safeguarding humid mountain forests and their connectivity to lowland forests to prevent an African biodiversity crisis.

## Data availability

The phylogenetic tree used in this study can be found on the GitHub repository of Joel Nitta at: https://fernphy.github.io/

The biogeographic distribution of the species of the phylogenetic tree can be found at: https://www.worldplants.de/world-ferns/ferns-and-lycophytes-list

Data used for the GLMM analysis belong to Anna Weigand and have been previously published here https://doi.org/10.1111/jbi.13782.

## Code availability

R code used to perform the ancestral range reconstruction analysis can be found at: GitHub - DMelladoMansilla/DispersalEventsReconstructionAfricanFerns

## Acknowledgments

MK acknowledges the financial support provided by the Swiss National Science Foundation (Grant No. SNF<u>310030L_146906</u>). The authors thank Dirk N. Karger for providing additional CHELSA layers. We also thank the valuable discussions provided by Sarah Noben, Mario Coiro, Merten Ehmig, Yanis Bouchenak-Khelladi, and Gabriel Ortega-Solis.

## Author contributions

AW, DMM, and MK designed the study, PW helped with the regional model, PW and HK provided regional data, AW and DMM performed the analyses. All authors contributed to the writing.

## Methods

### African fern species richness predictions

Information on 1243 regional entities with known fern species richness was compiled from the GIFT database ^51^. Using the programming environment R (version 3.2.2, R Core Team 2016), we extracted current environmental information (on a 1×1 km cell size) of all cells within each regional fern richness polygon, and calculated summary statistics (mean, median, 95%-quantile). Using generalized linear models, we tested several predictors to find the best model for global predictions. When necessary to improve linearity, predictors were log-transformed.

The final predictors were area, elevational range, potential evapotranspiration, mean annual cloud frequency ^32^, and habitat homogeneity (second order ^52^) as well as aridity index, temperature annual range, annual precipitation, and precipitation of warmest quarter ^53^.

Botanical continent (based on the continental scheme (level 1) of the Taxonomic Database Working Group ^54^ but differing in that the layer 2 division “Western Indian ocean” was separated from Africa for this study, Fig. 2) was included as a factor to allow differences in the magnitude of species richness for each continent. The model was projected onto a hexagonal equal-area grid with a grid cell size of 7,666 km2 ^55^. Grid cells with a landmass of less than one-quarter were excluded to avoid underestimating the richness of coastal or island regions.

To test whether current climate suitability is driving the lower species numbers in Africa, we trained our richness-climate-model on only Neotropical and Tropical Asian data (excluding Africa) and predicted it across all three regions. By comparing the resulting model with the original model based on all input data, we can identify the number of ferns that the African climate could support but are not observed. By comparing the species numbers predicted by the richness-climate-relationship trained on Neotropical and tropical Asian data, and the numbers resulting from the model also trained on African data, we estimated how many species the African climate should be able to support and how many we find (Fig. 1b). Additionally, we correlated this lack of species numbers with climatic predictors (Fig. 1c).

### Ancestral range reconstruction

To reconstruct the biogeographic history of fern lineages, we used the time-calibrated and continuously updated Fern Tree of Life (FTOL ^29^). This phylogeny includes 5,742 fern species and is the most comprehensive to date, with species richness representation varying across clades, ranging from 30% for Gleicheniales to 69% for Osmundales.

We classified the species in the FTOL into five biogeographical areas: Tropical America, Tropical Africa, Tropical Asia, Western Indian Ocean, and Extratropical regions (northern subtropical and arctic regions) (Fig. 2). Western Indian Ocean was separated from Africa due to its potential role as a stepping stone from Asia to Africa, which remains puzzling ^56^. The current distribution of the species in these five areas was obtained using the R package GIFT version 1.3.2 ^57^ and Hassler’s Checklist of Ferns and Lycophytes of the World (World Ferns) ^58^. The species richness in each area was as follows: 2091 species in Tropical America, 444 in Tropical Africa, 2784 in Tropical Asia, 496 in Western Indian ocean, and 1911 species in the Extratropical region. We estimated the representativeness of species richness for each area according to the information available on World Ferns for the overall species richness of each area, which varies from 50% for Tropical America, 66% for Tropical Africa, 54% for Tropical Asia, 71% for the Western Indian Ocean, and 86% for the Extratropical region.

We conducted an ancestral range reconstruction analysis using the R package BioGeoBEARS version 1.1.1^59^. First, we fitted the likelihood versions of the biogeographic models DEC (dispersal-extinction-cladogenesis ^60^) and its variant DEC*^61^ models, using the Akaike Information Criterion (AIC) to identify the model that best fitted our data. Both models allow a range of distributional processes suitable for widespread lineages, including anagenetic range evolution (i.e., dispersal, extirpation) and cladogenetic range evolution (i.e., narrow sympatry, subset sympatry, allopatry, and full sympatry)^60^. We excluded the founder-event versions due to the criticisms they have received^62^. We evaluated the other two commonly used biogeographical models, DIVALIKE^63^ and BAYAREALIKE^64^, but excluded them from the results as their assumptions do not align with the biogeography of ferns (Supplementary Table 2). Specifically, DIVALIKE does not permit the inheritance of sympatric subset areas, and BAYAREALIKE does not allow for the inheritance of cladogenetic areas, assuming no range evolution during cladogenesis^63–65^.

Since we were interested in the dynamics of specific periods in fern evolution, we used the time-stratified version of the models but with unconstrained, constant geography. The time periods used were: 421 to 66 mya, the Paleocene [66–56 mya], Eocene [56–33.9 mya], Oligocene [33.9–23.03 mya], Miocene [23.03–5.33 mya], Pliocene [5.33–2.58 mya], and Pleistocene [2.58–0 mya]. This matrix enabled us to obtain subsequent results on dispersal events in a more efficient and systematic way, while model fitting remained the same regardless of the inclusion of time-stratification (Supplementary Table 2). The best model, according to Akaike, DEC*, was used to generate 100 biogeographic stochastic mappings (BSM). BSM enables the generation of simulated histories or realizations with specific timings of biogeographic events across all branches of the phylogeny. We estimated the dispersal events, the number of lineages that survived in each area over the last 66 my, and the ancestral area of species currently inhabiting each area by calculating the mean and standard deviation of all events from the 100 BSMs. For long-term surviving lineages originating in more than one area, we counted for each continent separately into how many species the lineage diversified. All counts related to dispersal events are detailed in Table 3 of the Supplementary Information section.

Based on paleoclimatic maps based on lithologic indicators ^66^, we discuss possible processes behind different species numbers between continents and for extinctions and re-colonizations, specifically on the African continent.

## Supplementary Information

**Supplementary table 1.** Model comparison and predictor estimates for glm excluding (glm_base) and including (glm_botcont) botanical continent.

| Model | Predictor | Estimate | Std. Error | z value | Pr(>\|z\|) | AIC | R.squared |
| --- | --- | --- | --- | --- | --- | --- | --- |
| glm_base | (Intercept) | -6.123 | 0.061 | -100.811 | 0.000 *** | 54747 | 0.689 |
|  | log_area | 0.380 | 0.004 | 100.693 | 0.000 *** |  |  |
|  | log_q75_CHELSA_bio10_12 | 1.302 | 0.024 | 54.442 | 0.000 *** |  |  |
|  | log_q75_CHELSA_bio10_18 | 0.776 | 0.013 | 61.463 | 0.000 *** |  |  |
|  | log_range_elev | 0.818 | 0.009 | 90.408 | 0.000 *** |  |  |
|  | med_CHELSA_Aridity | -0.162 | 0.005 | -35.020 | 0.000 *** |  |  |
|  | med_CHELSA_bio10_7 | -0.008 | 0.001 | -12.820 | 0.000 *** |  |  |
|  | q75_Homogeneity_01_05_1km_uint16 | -0.689 | 0.033 | -20.926 | 0.000 *** |  |  |
|  | q75_MODCF_meanannual | 0.012 | 0.000 | 35.081 | 0.000 *** |  |  |
|  | q75_pet_he_yr | 0.000 | 0.000 | 25.500 | 0.000 *** |  |  |
| glm_botcont | Intercept) | -5.346 | 0.065 | -82.645 | 0.000 *** | 48501 | 0.745 |
|  | bot_cont_madaANTARCTIC | -0.510 | 0.068 | -7.459 | 0.000 *** |  |  |
|  | bot_cont_madaASIA-TEMPERATE | 1.128 | 0.018 | 64.067 | 0.000 *** |  |  |
|  | bot_cont_madaASIA-TROPICAL | 0.661 | 0.015 | 43.074 | 0.000 *** |  |  |
|  | bot_cont_madaAUSTRALASIA | 0.378 | 0.020 | 19.033 | 0.000 *** |  |  |
|  | bot_cont_madaEUROPE | 0.330 | 0.021 | 15.842 | 0.000 *** |  |  |
|  | bot_cont_madaMADAGASCAR | 0.497 | 0.029 | 16.865 | 0.000 *** |  |  |
|  | bot_cont_madaNORTHERN AMERICA | 0.737 | 0.016 | 44.753 | 0.000 *** |  |  |
|  | bot_cont_madaPACIFIC | 0.367 | 0.023 | 15.653 | 0.000 *** |  |  |
|  | bot_cont_madaSOUTHERN AMERICA | 0.415 | 0.014 | 30.069 | 0.000 *** |  |  |
|  | log_area | 0.391 | 0.004 | 100.105 | 0.000 *** |  |  |
|  | log_q75_CHELSA_bio10_12 | 1.190 | 0.024 | 49.435 | 0.000 *** |  |  |
|  | log_q75_CHELSA_bio10_18 | 0.707 | 0.012 | 57.584 | 0.000 *** |  |  |
|  | log_range_elev | 0.785 | 0.009 | 86.443 | 0.000 *** |  |  |
|  | med_CHELSA_Aridity | -0.207 | 0.005 | -42.983 | 0.000 *** |  |  |
|  | med_CHELSA_bio10_7 | -0.029 | 0.001 | -40.982 | 0.000 *** |  |  |
|  | q75_Homogeneity_01_05_1km_uint16 | -0.687 | 0.034 | -20.164 | 0.000 *** |  |  |
|  | q75_MODCF_meanannual | 0.010 | 0.000 | 25.596 | 0.000 *** |  |  |
|  | q75_pet_he_yr | 0.000 | 0.000 | 19.002 | 0.000 *** |  |  |

**Supplementary table 2.**
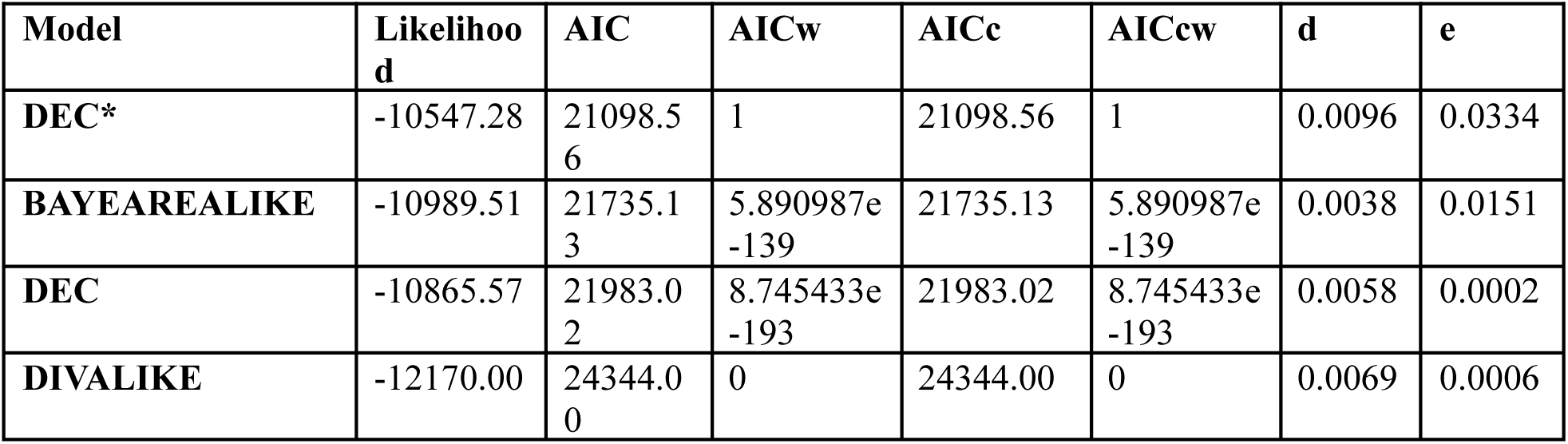
Likelihood versions of the models tested in this study. The table shows the likelihoods, Akaike values, and the dispersal rate (d) and extinction rate (e) parameters for each model.

**Supplementary table 3.** Means and standard deviations of all the dispersal events (from to) extracted from the 100 biogeographic stochastic mappings of the DEC* model. WIO: Western Indian ocean.

|  | **Extratropical** | **Tropical America** | **Tropical Asia** | **WIO** | **Tropical Africa** |
| --- | --- | --- | --- | --- | --- |
| **Extratropical** | 0 | 254.2 ± 14.6 | 482.2 ± 24.8 | 121.7 ± 11.3 | 133.6 ± 11.2 |
| **Tropical America** | 331.3 ± 18 | 0 | 186.2 ± 14 | 164.7 ± 12.7 | 164.3 ± 12.6 |
| **Tropical Asia** | 847.8 ± 26.2 | 247.6 ± 15.1 | 0 | 231.2 ± 13.4 | 224.3 ± 12.8 |
| **WIO** | 91.4 ± 9.6 | 85.0 ± 9.3 | 91.2 ± 10 | 0 | 137.4 ± 11 |
| **Tropical Africa** | 95.5 ± 10.2 | 88.3 ± 9.9 | 96.8 ± 11.2 | 151.4 ± 10.6 | 0 |

